# AxonDeep: Automated Optic Nerve Axon Segmentation in Mice with Deep Learning

**DOI:** 10.1101/2021.05.21.445196

**Authors:** Wenxiang Deng, Adam Hedberg-Buenz, Dana A. Soukup, Sima Taghizadeh, Kai Wang, Michael G. Anderson, Mona K. Garvin

**Author notes:** Corresponding author: Mona K. Garvin, Ph.D., 4318 Seamans Center for the Engineering Arts and Sciences, The University of Iowa, Iowa City, IA 52242. Contributed equally (in first-author role) (Wenxiang Deng, Adam Hedberg-Buenz). Contributed equally (in senior-author role) (Michael G. Anderson, Mona K. Garvin). Disclosure: **W. Deng**, None; **A. Hedberg-Buenz**, None; **D.A. Soukup**, None; **S. Taghizadeh**, None; **K. Wang**, None; **M.G. Anderson,** None, **M.K. Garvin**, None.

## Abstract

**Purpose:** Optic nerve damage is the principal feature of glaucoma and contributes to vision loss in many diseases. In animal models, nerve health has traditionally been assessed by human experts that grade damage qualitatively or manually quantify axons from sampling limited areas from histologic cross sections of nerve. Both approaches are prone to variability and are time consuming. First-generation automated approaches have begun to emerge, but all have significant shortcomings. Here, we seek improvements through use of deep-learning approaches for segmenting and quantifying axons from cross sections of mouse optic nerve.

**Methods:** Two deep-learning approaches were developed and evaluated: (1) a traditional supervised approach using a fully convolutional network trained with only labeled data and (2) a semi-supervised approach trained with both labeled and unlabeled data using a generative-adversarial-network framework.

**Results:** From comparisons with an independent test set of images with manually marked axon centers and boundaries, both deep-learning approaches outperformed an existing baseline automated approach and similarly to two independent experts. Performance of the semi-supervised approach was superior and implemented into AxonDeep.

**Conclusion:** AxonDeep performs automated quantification and segmentation of axons from healthy appearing nerves, and those with mild to moderate degrees of damage, similar to that of experts without the variability and constraints associated with manual performance.

**Translational Relevance:** Use of deep learning for axon quantification provides rapid, objective, and higher throughput analysis of optic nerve that would otherwise not be possible.

## Introduction

Retinal ganglion cell (RGC) loss is the primary feature of glaucoma, a leading cause of irreversible vision loss.^1, 2^ RGCs are also damaged as a part of several other diseases, including forms of traumatic brain injury (TBI),^3, 4^ diabetes,^5^ multiple sclerosis,^6, 7^ and Alzheimer’s disease,^8^ among others.^9^ To evaluate RGC damage, the two fundamental options are to quantify RGC soma in the retina or RGC axons in the optic nerve, which in health typically have a 1:1 relationship. Because axon damage can occur earlier than somal loss,^10, 11^ there are many situations in which it is useful to quantify both. While the identification of RGC-specific markers has helped advance techniques for quantification of RGC soma,^12, 13^ techniques for quantification of RGC axons, which are smaller and more difficult to label, have lagged. Manual counting of axons has traditionally been adopted by human experts.^14–18^ However, manually counting is not only time-consuming, but prone to vary between different experts. Grading of optic nerve appearance based on pathological features has been useful for making some advances,^19–24^ but is even more subjective and prone to variation.

First-generation automated approaches for quantifying axons, such as AxonJ^25^, AxonMaster^26, 27^, AxoNet^28^, and use of QuPath^29^ have several known limitations. For example, AxonJ, which members of our group were involved in developing, was designed to recognize axons only in healthy, not diseased, optic nerves^25^. AxonJ has been demonstrated to not perform well with damaged nerves^29^. Likewise, AxonMaster was designed using healthy, not diseased, nerve tissue. It is also relevant that few of the existing approaches were developed using mice (AxonMaster, non-human primates; AxoNet, rats; QuPath, rats). It can be expected that deep-learning approaches, especially given their success in numerous medical application domains^30–35^ ultimately would be more robust. In fact, recently, the deep-learning approach AxoNet^28^ has been proposed for providing axon counts; however, this approach still does not provide direct segmentation of the axons and thus the ability to compute additional potential quantitative measures (such as area distributions) using this approach is limited.

In the present work, we propose a deep-learning approach, named AxonDeep, for the segmentation of optic nerve axons in murine tissue. This work was motivated by the need to overcome some of the known limitations of first-generation approaches^25^ and a desire to obtain complete segmentations (allowing quantitative measures of multiple axon measurements, beyond just counts) that are not possible with the current tools.^28^ Our deep-learning architecture is based on recent work on asymmetric network structures whereby a deep encoder combined with a light-weight decoder can provide an improved performance over symmetric deep-learning architectures for image segmentation problems with more complex scenes.^23, 27^ In developing AxonDeep, two deep-learning approaches were evaluated: (1) a traditional supervised approach using a fully convolutional network (FCN) trained with only labeled data and (2) a semi-supervised approach trained with both labeled and unlabeled data using a generative-adversarial-network framework. Utilizing the semi-supervised approach to be able to also take advantage of unlabeled data, in addition to labeled data, during training was motivated by a need to help address the challenges associated with the manual effort required to generate training sets. Both deep-learning approaches were compared to the AxonJ approach, with the semi-supervised approach found to have the better overall performance. Thus, we introduce AxonDeep as a deep-learning-based axon segmentation tool trained based upon a semi-supervised approach.

## Methods

### Procurement and preparation of mouse optic nerve specimens for image analysis

*Mice*. Optic nerves (*n=*56 nerves, 1 nerve from each of 56 mice) were collected from three different mouse strains modeling various presentations of healthy and diseased optic nerves: DBA/2J with various degrees of an inherited age-related form of glaucoma (*n=*11 nerves);^36, 37^ D2.B6-*Lyst^bg-J^*/Andm (abbreviated hereafter as D2.*Lyst*) with healthy optic nerves (*n*=5 nerves); C57BL/6J that had been subjected to either blast-induced traumatic brain injury (TBI blast) within an enclosed chamber or sham treatment (TBI sham; *n=*27 nerves);^3, 4, 22^ and Diversity Outbred (J:DO) with healthy optic nerves but predicted to exhibit genetic background-dependent natural variability in optic nerve features (*n=*13 nerves).^38, 39^ Tissues in the current study were collected from mice contributing to prior publications,^40, 41^ however all data reported herein arise from new analyses performed uniquely for this study. All mice used in this study were originally purchased from The Jackson Laboratory (Bar Harbor, ME). All animals were treated in accordance with the ARVO Statement for the Use of Animals in Ophthalmic and Vision Research. All experimental protocols were approved by the Institutional Animal Care and Use Committee of the University of Iowa.

#### Nerve processing

Nerves were processed for histology as previously described.^17, 25, 42^ In brief, mice were euthanized by carbon dioxide inhalation with subsequent decapitation. Heads were collected and the skulls opened before fixation in half-strength Karnovsky’s fixative (2% paraformaldehyde, 2.5% glutaraldehyde in 0.1 M sodium cacodylate) at 4°C for 16 hours. Optic nerves were dissected from brains and drop fixed in the same fixative for an additional 16 hours at 4°C. Nerves were stained with 1% osmium tetroxide, dehydrated in graded acetone (30%-100%), infiltrated and embedded in resin (Eponate-12; Ted Pella, Redding, CA), and polymerized in a 65°C oven. Semithin (1-µm) cross sections were cut, transferred to glass slides, stained with 1% paraphenylenediamine (PPD), and mounted.

### Procurement of images for training, validation, and testing of the network

Light micrographs (physical dimensions: 90.2 x 67.5 µm; resolution: 4140 x 3096 px) were collected from stained optic nerve cross sections at a total magnification of 1000X using identical camera settings, as previously described.^17, 25, 42^ In brief, light micrographs were acquired from representative (i.e., a field not atypical from the rest of the nerve) and non-overlapping fields from one cross-section of each optic nerve using an upright light microscope (BX52, Olympus) equipped with a CCD camera (DP-72, Olympus).

#### Separation of images for purposes of training/validation and testing of the deep-learning networks

Optic nerve images were divided into two major sub-divisions used for different purposes, a “training/validation” set and a “testing” set (Figure 1). To ensure an equal distribution of nerves with comparable levels of damage between the two sets, a qualitative damage grade was first assigned to each nerve by consensus among a panel of three independent graders (1 = mild or no damage, 2 = moderate damage, and 3 = severe damage), as previously described.^14^ Representative examples of each damage grade are shown in S. Figure 1. Within grade-1 nerves, two subgroups were utilized: i) grade-1 no apparent damage, defined as nerves from strains of healthy mice and histologically free of damage, and ii) grade-1 mild damage from disease models (i.e. DBA/2J^36, 43^ or blast-induced TBI^4, 44^) at early stages in which no damage was yet apparent, but could be present sub-clinically. Among the 56 nerves, images from 38 nerves (1-2 images per nerve from 38 mice) were assigned to the training/validation set (24 grade-1 nerves: 12 with grade-1 no apparent damage and 12 with grade-1 mild damage; 12 grade-2 nerves; and 2 grade-3 nerves) and images from the remaining 18 nerves (1 image per nerve from of 18 mice) were assigned to the test set (12 grade-1 nerves: 6 with grade-1 no apparent damage and 6 with grade-1 mild damage; 6 grade-2 nerves). Axon number in mice is highly dependent on genetic background^45^; therefore, the grade-1 C57BL/6J, grade-1 DBA/2J, and every J:DO nerve is expected to have a different number of axons. Because of potential challenges in segmenting (manually or automatically) severely damaged nerves, the two grade-3 nerves were assigned to the training set (and only used as additional unlabeled examples in the semi-supervised approach). Of the challenges with segmenting grade-3 nerves, some relate to the fibrotic and gliotic changes that drastically alter gross nerve appearance. The current study was designed to emphasize training AxonDeep to recognize axons from nerves with more of a continuum in gross appearances (normal, mild damage, moderate damage) and limited testing to only grade-1 and grade-2 nerves. The remaining 36 mild and moderate nerves in the combined training/validation set were further randomly divided into 26 nerves to be used for training and 10 nerves to be used for validation. Note that, as is standard practice with deep-learning techniques, the training process was used for automatically determining the trainable network weights, whereas the validation process was used for deciding hyperparameters and tuning the network. The random division of the 36 mild/moderate nerves into training and validation sets resulted in 17 grade-1 nerves and 9 grade-2 nerves in the training set, and 7 grade-1 nerves and 3 grade-2 nerves with moderate damage in the validation set. A reference segmentation was manually obtained on a 1024x1024 sub-field as described in the next section on each of the 36 mild/moderate nerves allocated to the training/validation sets (6762 total axons on the 26 training images and 2668 total axons on the 10 validation images); however, all 50 available 4140x3096 full-sized images (in the training set) were also used as unlabeled images (with more than 150,000 axons total) for helping to train the semi-supervised approach (see semi-supervised approach).

**Figure 1.**
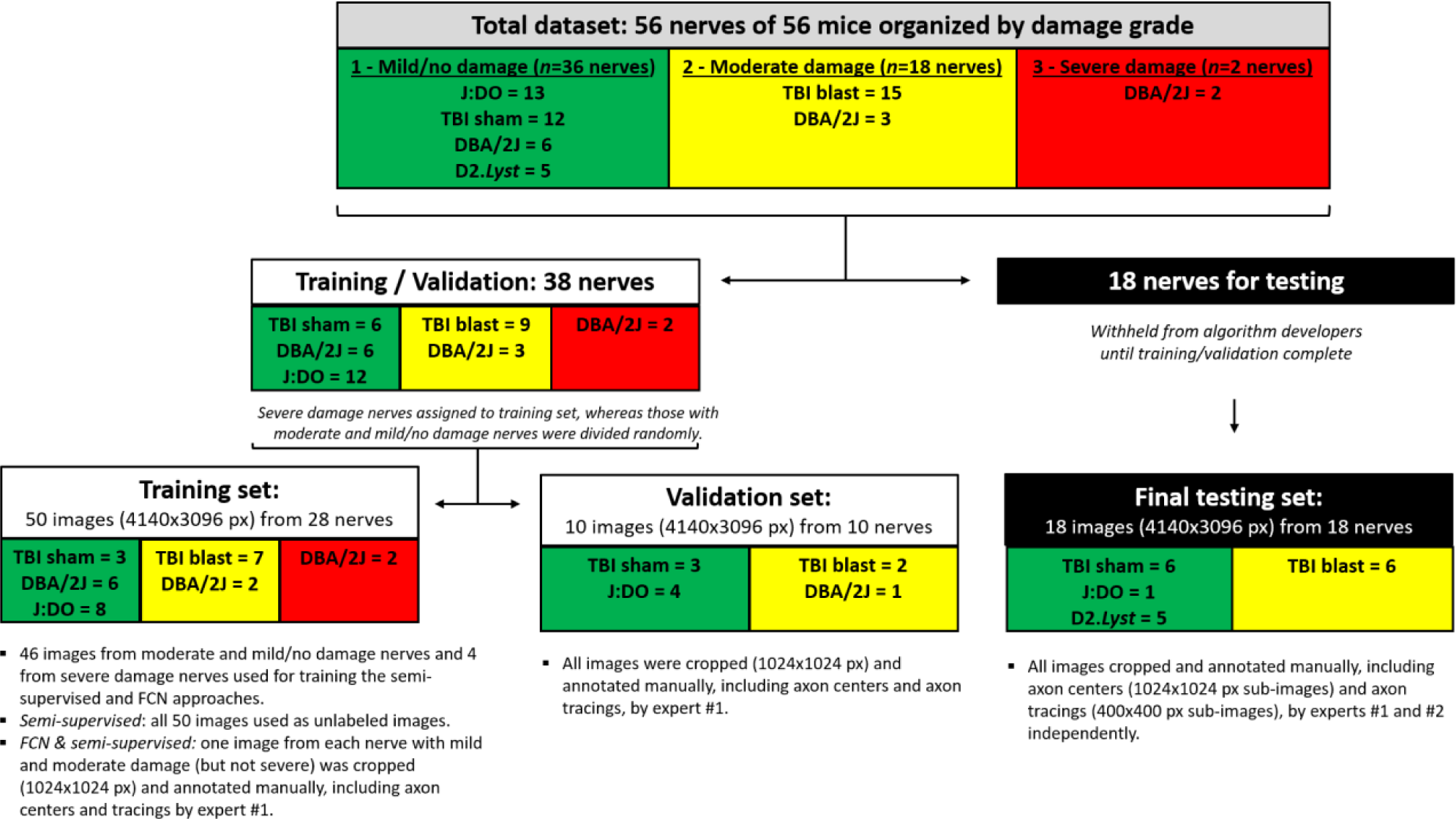
Flowchart of datasets, experimental design, and progression of tool development. The total dataset was composed of a diverse collection of optic nerve specimens from multiple genotypes and strains with natural phenotypic variability (i.e. J:DO), normal health (i.e. D2.Lyst) and forms of damage resulting from naturally occurring disease (i.e. DBA/2J) or inducible injury (i.e. TBI blast). Nerves were qualitatively graded (grade 1: mild/no damage [green]; grade 2: moderate damage [yellow]; grade 3: severe damage [red]) and divided into cohorts with 28 nerves for training, 10 nerves for validation, and 18 nerves for final testing. Across each set, the composition of nerves by damage grade remained consistent for the validation and training sets (ratio of [2:1] grade 1:grade 2 nerve); note that the tool was not designed to quantitate grade 3 nerves with severe damage and that only unlabeled images as part of the training set included grade 3 nerves. Based on annotations by expert #1, a total of: 6,762 axons centers were marked in the training set, 2,668 axons centers were marked in the validation set, 3,317 axons centers were marked and 1,103 axons were traced in the final testing set. From the un-annotated set of images (n=50) used in the training set, there were in excess of 150,000 axons.

### Obtaining reference axon segmentations to be used for training and validation

As our deep-learning networks provide pixel-based marking of axons, our reference standard to train the network and optimize hyperparameters needed to include complete segmentations of the axons. In other words, we needed to obtain binary axon masks (white = axon pixels; black = non-axon pixels) alongside the original images to train the networks. The supervised network required all input images for training to have complete segmentation information and the semi-supervised network still required complete segmentation information for a subset of images. Because obtaining completely traced axons from scratch is labor intensive (even for obtaining the subset of complete segmentations needed for the semi-supervised approach), for training purposes, our strategy to obtain complete segmentations involved manually marking axon centers in combination with manually correcting the boundaries of an automated segmentation. (Note that, as discussed later, our strategy for obtaining complete tracings for evaluation purposes on the test set did not involve an automated step, as was used in the training stage, to avoid any bias associated with involving an automated segmentation.)

More specifically, obtaining the reference tracings to be used for training involved the following. From the 4140x3096 image(s) available per nerve for training purposes, we first randomly selected one of the images and cropped a sub-field of size 1024x1024 (location selected at random) for purposes of obtaining complete segmentation tracings. On each of these 1024x1024 sub-fields, we ran the baseline AxonJ approach followed by additional morphological smoothing processes to obtain a smooth starting segmentation for purposes of manual editing. Figure 2(B) shows an example segmentation of the axon image in Figure 2(A). In addition, we independently obtained manual center-point markings by placing a center point at an approximate center of each axon (dead/dying axons are marked separately, but not used in this work), as shown in Figure 2(C).

**Figure 2.**
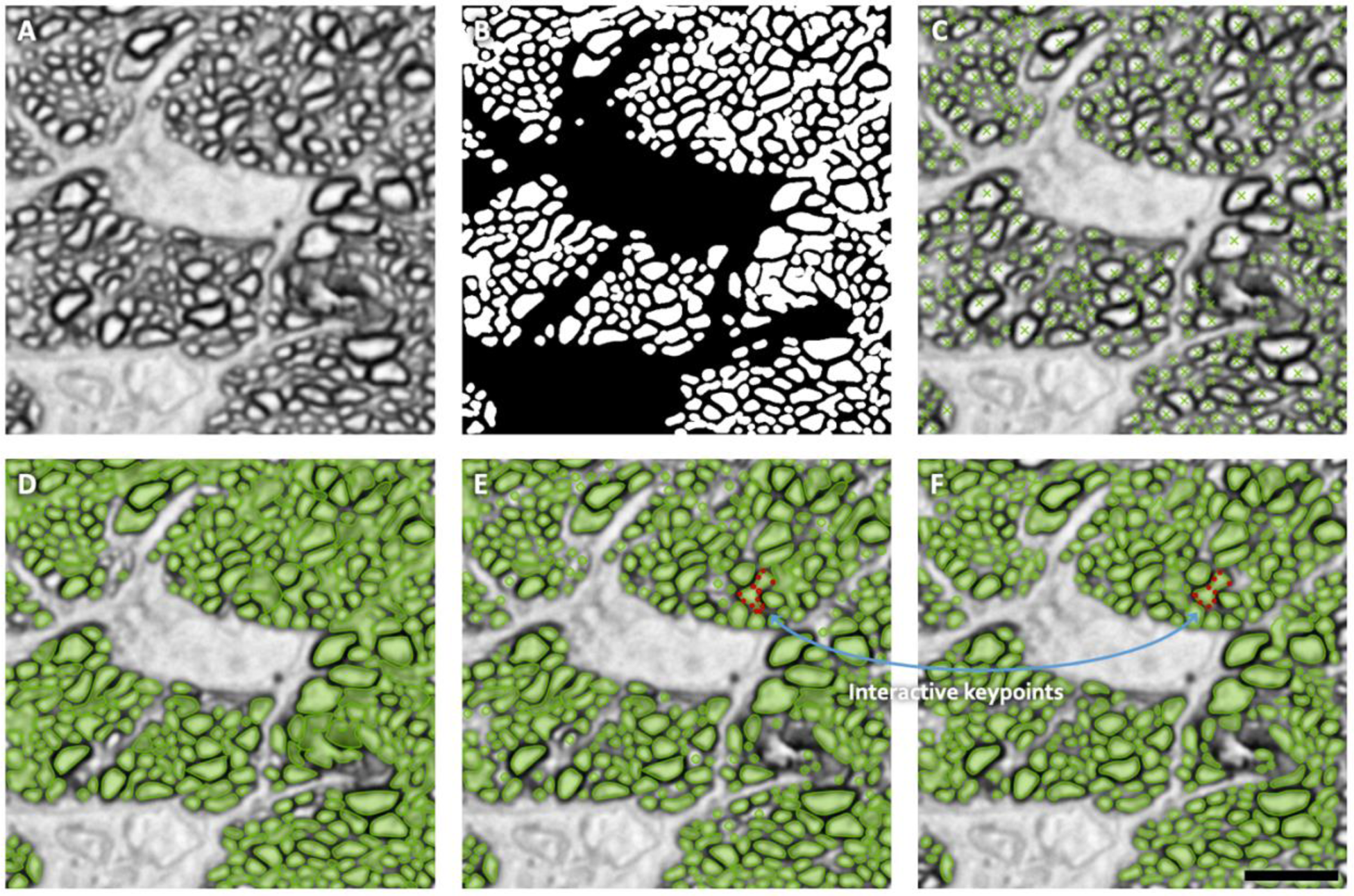
The coupling of manually corrected automated segmentations with manual tracing of center marks of axons was used to construct the reference segmentations for training. (A) Light micrograph of a paraphenylene diamine-stained optic nerve cross section in enhanced format (histogram equalized for better visualization) used as an input image for training. (B) Binary AxonJ result. (C) Manually marked axon centers. (D) AxonJ result in (B) displayed using interactive GUI (clicking on an axon would cause interactive keypoints to appear). (E) Pruned AxonJ result with red interactive keypoints for a sample axon contour indicated. (F) Result of editing sample axon contour. By combining automated segmentation (B), manually traced center marks (C), and manual corrections (E)(F), references for training data can be obtained. Scale bar = 5 µm.

Next, the border of each segmented axon was detected,^46, 47^ and the contours were converted into representative keypoints. Here we used a simple heuristic approach to generate the points. More specifically, the Ramer Douglas Peucker (RDP) algorithm^48^ was used to generate a keypoint representation. This greedy algorithm iteratively found a polyline that is close to the contour we segmented, with the maximum distance from the interpolated line between keypoints and the true edge was less than a predetermined distance ε. Here we used an ε of 5.0 pixels. These keypoints were then visualized with a GUI interface, as shown in Figure 2(D).

We next combined the results from the manually marked centers (Figure 2(C)) and smoothed AxonJ contours as follows: 1. False positives (i.e., axons segmented by AxonJ, but not marked by the human expert) were excluded from the mappings; 2. false negatives (i.e., axons not segmented by AxonJ) were annotated with a small starting circle. An example pruned map can be seen in Figure 2(E). Finally, the contours were manually corrected by changing the locations (marked in red) of the interactive keypoints shown in Figure 2(E) and (F).

### Obtaining reference axon counts and segmentations to be used for final evaluation (testing)

For the final one-time evaluation of each trained approach on the test set, we also obtained reference manual axon counts and complete segmentations. However, unlike in training, in obtaining complete segmentations we traced the boundaries of the axons from scratch (i.e. not editing an automated result) to avoid any bias associated with use of an automated approach. More specifically, for each of the test images, after obtaining a random 1024x1024 crop from a raw full-sized image, as shown in Figure 3(A-B), approximated axon centers within the bounding box were manually marked to evaluate the axon count predictions (Figure 3C). The axons close (less than 120 pixels) to image borders were ignored in the evaluation because it is hard to accurately assess the axons when part of them are outside the border. For the pixel-based evaluation of the axon segmentations, a randomly chosen 400×400 sized cropped sub-field was used for complete tracing by the same expert (rather than editing an existing segmentation as was done for training). Figure 3(D) shows an example selection of the traced area in the bounding box. Note that a description of the metrics used for comparing the reference standard to the automated approaches for final evaluation appears in the Evaluation subsection.

**Figure 3.**
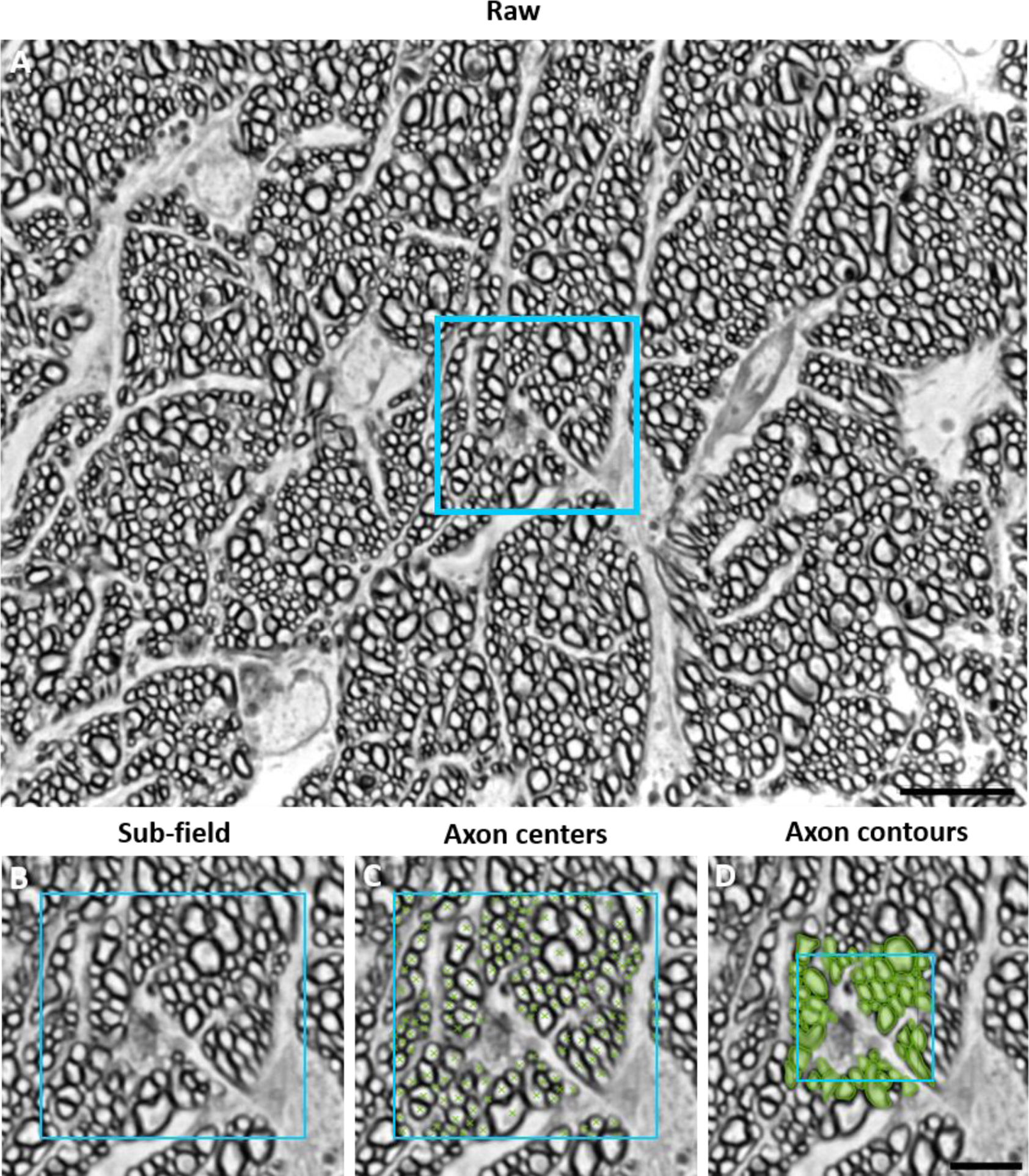
Manual annotation of axon centers and contours for evaluating tool performance. An example of progressive annotations on the same microscopic field from a paraphenylene diamine-stained optic nerve cross section in (A) raw full-sized form (4140x3096 px) and cropped sub-fields (1024x1024 px) (B) before manual annotation of (C) axon center marks (green x’s) and (D) axon tracings (green outlines and infilling; 400x400 px) in a smaller sub-field to provide a reference for axon counts and contours for final evaluation in the test set. Inset blue box denotes the same border for inclusion of axons for counting and tracing and elimination of edge effects for panels (A-D). Scale bar = 10 µm (A) and 5 µm (B-D).

### Deep-learning approach 1: Fully convolutional network (FCN) for segmentation

Figure 4 provides an illustration of the architecture of our FCN used to provide a pixel-level segmentation of the axons in the image. Overall, it uses an encoder-decoder framework with use of an established encoder (a deep ResNeXt-50^49^ in our case) in combination with a more lightweight, asymmetric (compared to the encoder) decoder. This type of architecture is an example of a feature pyramid network (FPN) and has been shown to be successful in image-segmentation and detection tasks.^23, 50–52^ For the encoder of the network, we used a deep ResNeXt-50^49^ architecture, which uses a very similar structure as ResNet^53^ while offering better performance. Use of a deep encoder was motivated by the need to consider the context of a relatively large surrounding region in determining whether a pixel belongs to an axon. Note the asymmetry of the encoder-decoder is different from the symmetric structure used in the popular U-Net framework popular in many pixel-based medical segmentation tasks.^23^

**Figure 4.**
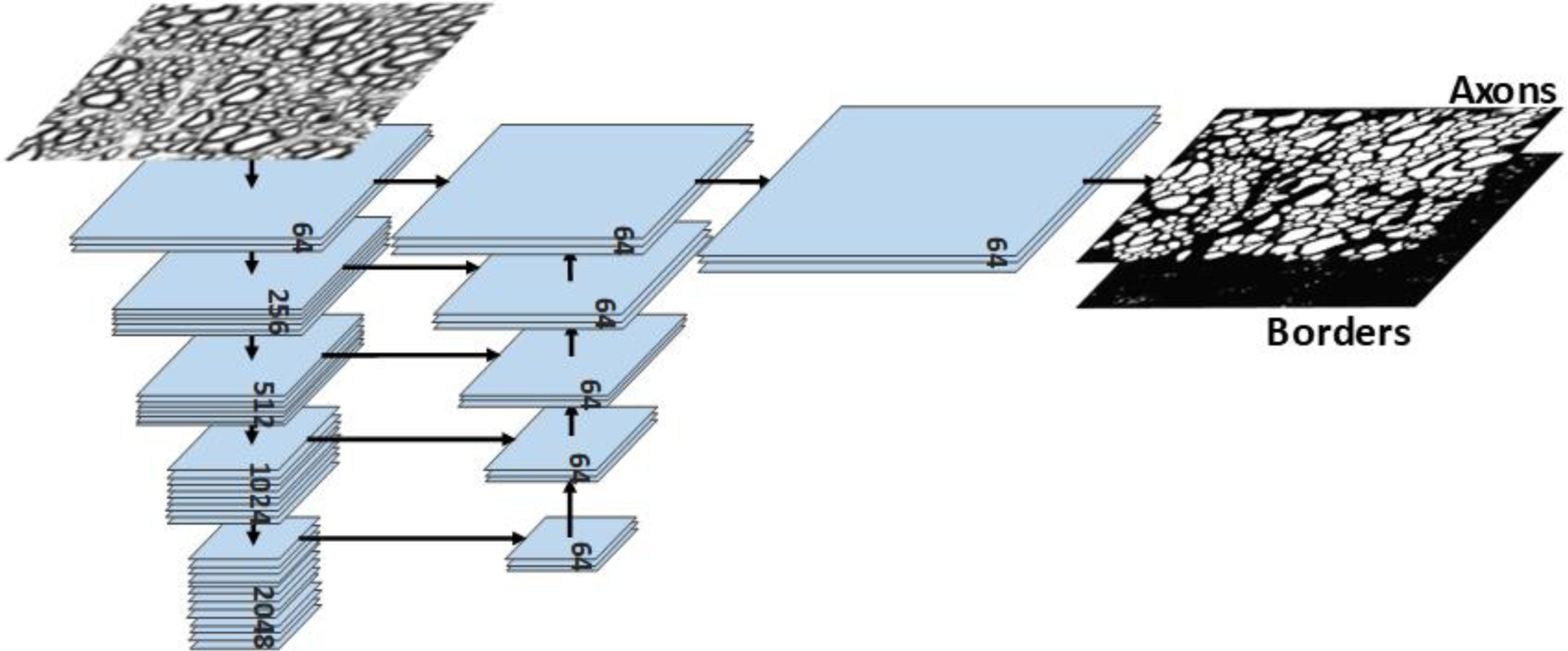
An illustration of the network for our FCN approach. The backbone network is paired with a light-weight feature pyramid (FPN) like decoder. The network takes a single channel axon image as input and outputs an axon probability map and borders between adjacent axons.

For the training data, in order to minimize the chance of adjacent axons being segmented as a single object, we separately predicted the axons and the borders between adjacent axons with two channels as output (as shown in Figure 4 and Figure 5(B)(C)). To generate the borders for training, we performed a morphological dilation on each manually corrected binary axon segmentation and found its overlap with the rest of the morphologically dilated axons. This overlap was defined as the border between adjacent axons (shown in Figure 5(C)). Note that at test time, these segmented borders can be used to better separate attached axon segmentation masks. As a result, a single channel of axon image with resolution divisible by 32 was used as input, and the network output consisted of two channels: an axon mask and the borders between nearby axons.

**Figure 5.**
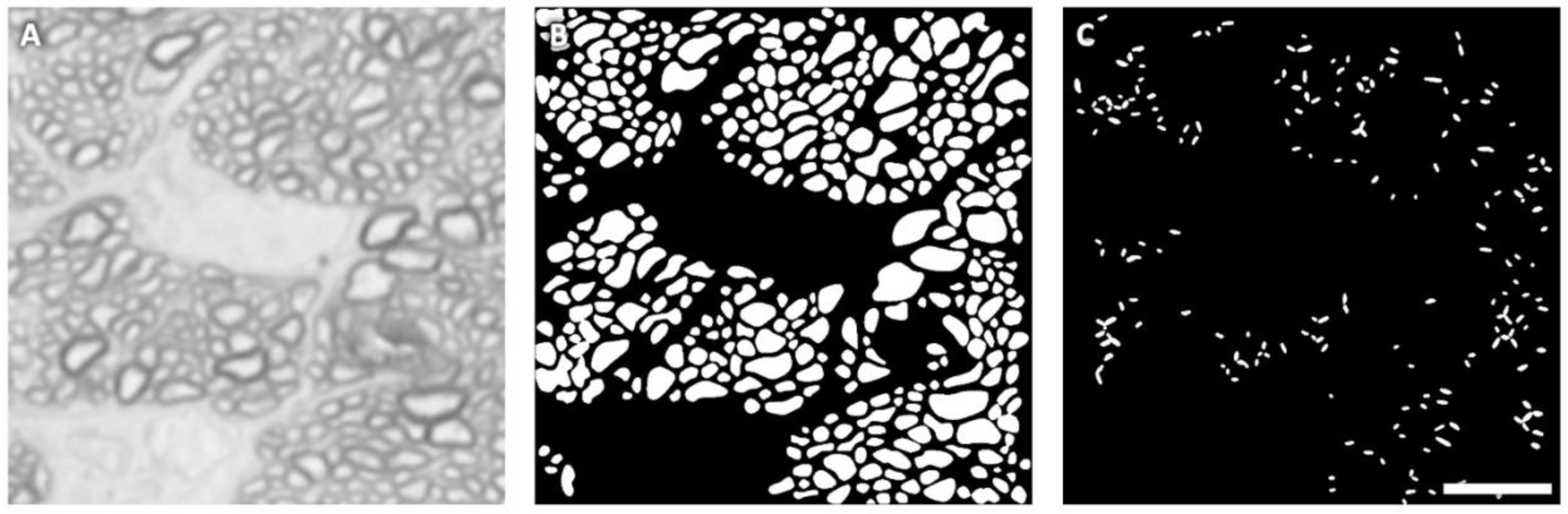
Examples of training data from analyses of optic nerve micrographs. An example of a (A) raw optic nerve image input, (B) generation of an axon mask (axons appear as white, whereas non-axons as black), and (C) borders between adjacent axons. Scale bar = 5 µm.

In order to train the network, similar to an approach used previously ^54^, at each training step, the total loss was a function combining a soft Dice loss (*L*_*Dics*_) and binary cross entropy loss (*L*_*BCE*_):

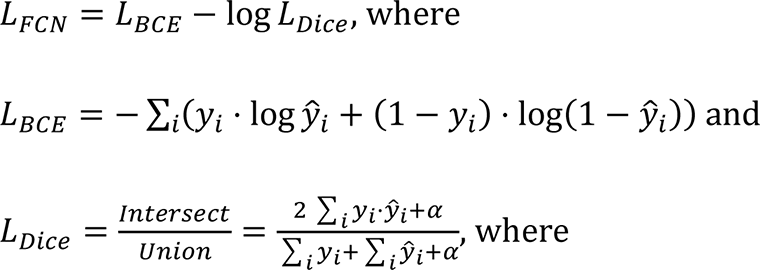

*y*_*i*_ and 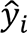 are corresponding truth and prediction at each pixel location *i*. Here α = 1 provides numerical stability and prevents cases of zeros.

At inference time, to generate the final output of the axon segmentation, a watershed algorithm^55^ was applied with axon segmentations minus adjacent borders as basins and axon segmentations as masks.^56^

### Deep-learning approach 2: Semi-supervised learning

To be able to take advantage of both labeled (time-consuming to acquire) and unlabeled data (i.e., just the input images without any manual annotations) during training, we also used a semi-supervised learning approach based on incorporating the FCN architecture described above into a generative-adversarial-network (GAN) setup. In a traditional GAN,^57^ a generator subnetwork *G* (designed to generate realistic images from noise) is simultaneously trained with a discriminator network *D* (designed to differentiate between real images and those generated by the generator). Because the subnetworks are trained together (in an alternating fashion), the networks “compete” in a minimax game so that ultimately the generator subnetwork can generate realistic images on its own (in order to fool the discriminator to try to win the competition). In other words, the addition of the discriminator subnetwork is used to help in the *training* of the generator subnetwork, but the discriminator subnetwork is not needed once training is complete. (Note that in other applications, one can actually keep the discriminator rather than the generator for purposes of having a starting point for a semi-supervised classifier as in Salimans et al.^58^ and take advantage of both labeled and unlabeled data for image-level *classification* tasks.) However, in our case, rather than just generating realistic-looking images, we wish to take input images and produce a resulting segmentation. Thus, we follow a similar approach as previously used^59, 60^ and a GAN-like framework to help train a segmentation network with both labeled and unlabeled data. An illustration of our approach can be seen in Figure 6. Intuitively, instead of having a generator *G* to take noise and generate realistic-looking images, we use a subnetwork *G* to take an input image and produce a resulting segmentation and a subnetwork *D* to differentiate (at a pixel level) between segmentations produced by the network *G* and reference segmentations (i.e., being able to tell if the segmentation, at each pixel, was from the generator subnetwork or used as the reference ground-truth). Here both *G* and *D* for our semi-supervised learning approach use the same network structure following the FCN approach in the previous section. Note that after training is complete, as with a traditional GAN, we can just use the trained *G* subnetwork (and not use the discriminator subnetwork) in practice (e.g., during testing) so that the inputs/outputs are just like they would be with the FCN described in the prior section.

**Figure 6.**
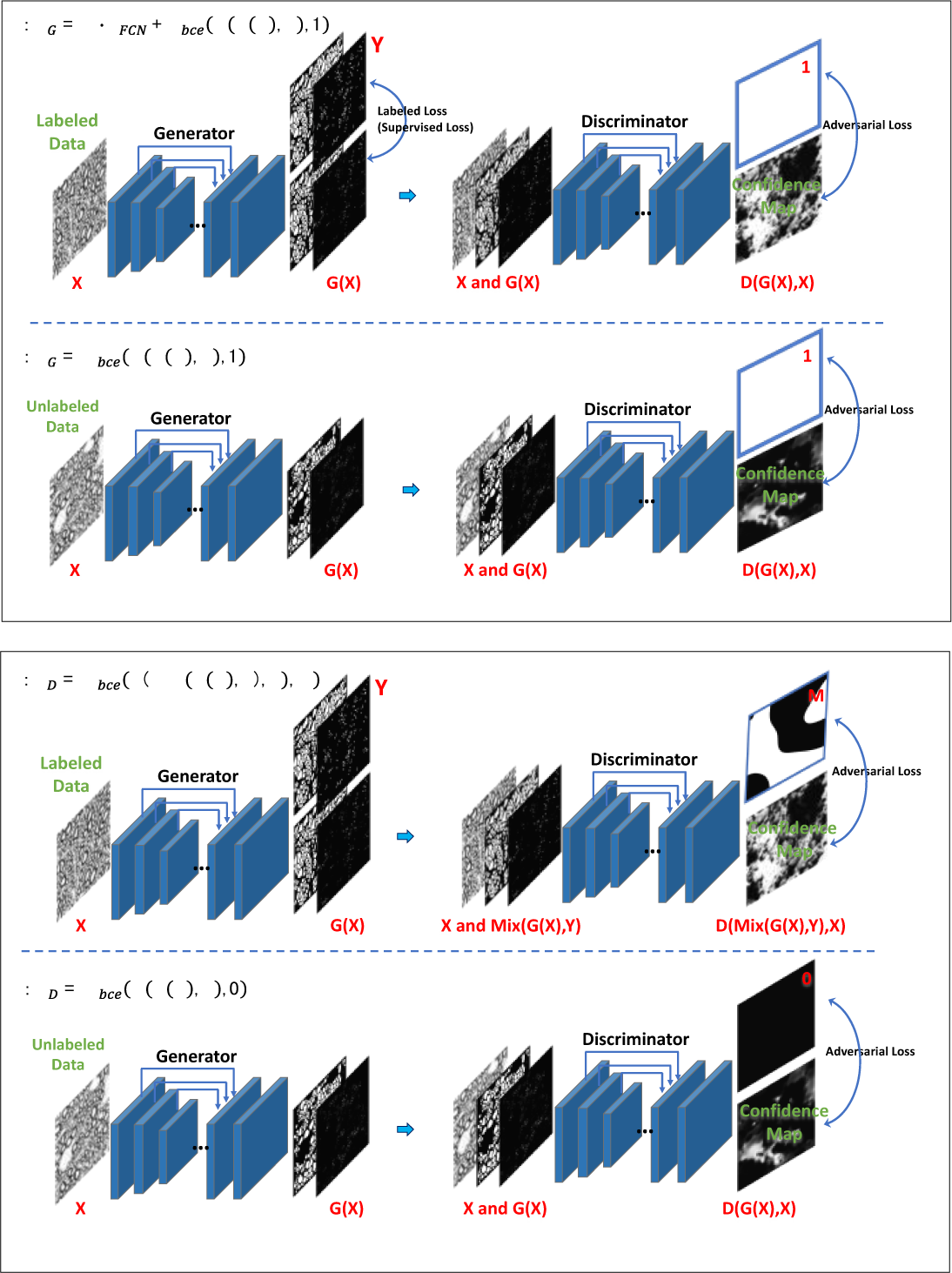
Training the semi-supervised approach alternates between updating the generator network weights and updating the discriminator network weights. Both the generator and discriminator have the same basic underlying architecture as the FCN approach. (After training only the generator network is retained as the final segmentation network.) The input to the generator is a raw axon image and the input to the discriminator is the raw axon image plus a segmentation output. The network uses both labeled data (with both axon images (X) and reference segmentations (Y) available) and unlabeled data (X) in updating the weights. (A) In updating the generator weights with labeled data, the loss function encourages the generator output to match the reference segmentation and to “fool” the discriminator in thinking that the generated output is a reference output. (B) In updating the generator weights with unlabeled data, the loss function encourages the generator to “fool” the discriminator in thinking that the generated output is a reference output. (C) In updating the discriminator weights with labeled data, the loss function encourages the discriminator to correctly output a 1 for pixels coming from a reference segmentation and a 0 for pixels coming from a generated segmentation. Note that a randomly mixed segmentation image (between the generated segmentation and reference segmentation) is provided as part of the input to the discriminator. (D) In updating the discriminator weights with unlabeled data, the loss function encourages the discriminator to correctly output a 0 for pixels not coming from a reference segmentation.

Just as with a traditional GAN, each training iteration alternates between training the *G* network and training the *D* network. When training the *G* network, the weights in the *D* network are frozen and won’t be updated (and vice-versa). Both labeled and unlabeled images can be used to update the weights (in each epoch, all 26 labeled images in the training set are used and a random subset of 26 of the 50 unlabeled images, to match the number of labeled images, is used). In training *G* (i.e., the segmentation network), when the input is a labeled image (i.e., the input *X* has a corresponding reference segmentation *Y*), the loss function is a combination of the same supervised loss used in the fully supervised approach as well as a binary cross-entropy loss based on passing the result of the generator (i.e., segmentation) network through the current version of the discriminator:

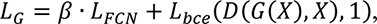

where *β* = 10 to balance supervised loss and GAN loss and *L*_*bce*_ is *L*_*BCE*_ with a plain Sigmoid function:

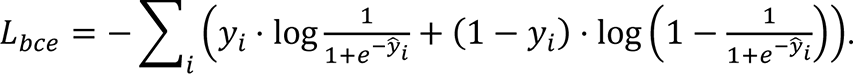

Note that in this case, the generator wants the discriminator to output a value of 1 at each pixel to indicate that the discriminator is fooled into thinking the generator’s output is a true manual tracing. When the input is an unlabeled image *X*, a reference labeled image is not available, so the loss function is solely based on the binary cross-entropy loss indicated in the second part of the equation above:

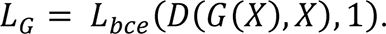

Similarly, in training *D* (the discriminator that tries to differentiate, on a pixel-by-pixel level, using both the segmentation map and original input image as input, whether the resulting segmentation came from the generator/segmentation network or a manual reference), when the input is an unlabeled image *X*, the loss function is also based on a binary cross-entropy loss, but this time to encourage the generator’s output to result in a value of 0 at each pixel location from the discriminator (i.e., the discriminator wants to correctly predict that the generator’s output is not a reference segmentation):

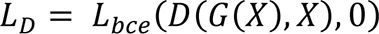

In training *D*, when the input is a labeled image (i.e., the input *X* has a corresponding reference segmentation *Y*), the current version of the generator is used to create the generated segmentation map, *G(X)*. A “mixed” image *Mix(G(X), Y)* is then created by randomly selecting each pixel to have a value either from the generated segmentation map *G(X)* or the reference image *Y*. A mask image *M* is created with values of 1 at locations where the pixels came from *Y* and 0 where pixels came from *G(X)*. The loss function used is the binary cross-entropy loss between the discriminator output (using the original image and mixed image as input) and the mask image of 1’s and 0’s:

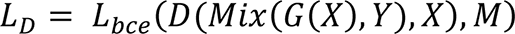

As before, this loss function will encourage the discriminator to have an output of 0 at locations where the input was from the generator network and an output of 1 at locations where the input was from the actual reference segmentation.

### Training strategies

To implement our approaches, the PyTorch framework was used for all of the experiments. A single Nvidia GeForce 1080Ti GPU with 12GB memory was used for training and testing. To train the networks, we used mini-batch stochastic gradient descent (Adam) as the optimizer.^61, 62^ The input image size for training was set to be 512×512, with a batch size of 4, randomly cropped from training images. Before each training mini-batch, data augmentation was applied to the input images with random resize, cropping, flip, Gaussian blur, and contrast changes.

For the learning rate, the FCN approach used an initial value of 1e-3 and was divided by a factor of 10 when the training loss plateaued. For the semi-supervised approach, a learning rate of 1e-4 for the *D* network and 1e-3 for the *G* network was used.

In order to determine when to stop training, the networks were evaluated on the validation set every 10 epochs. To balance the quality of axon segmentation and the count of axons generated from post-processing, we used an equal combination of accuracy and the absolute axon number difference between prediction and reference as the metric for the segmentation. More specifically, the receiver operating characteristic (ROC) curves with respect to prediction thresholds were measured and area under curve (AUC) of ROC was calculated as a measure of pixel accuracy. The network parameters providing the highest performance on the validation set was used as our final approach for evaluations.

### Evaluation

The AxonJ approach and our trained FCN and semi-supervised approach were used to provide both axon counts and complete segmentations for each of the 18 test images (corresponding to 3317 marked axon centers and 1103 fully traced axons from the first expert). The axon counts and complete tracings (obtained as discussed previously) from a single expert were used as the reference for comparisons with the automated approaches. The absolute percent axon count difference (as a percentage of the reference count, i.e., the absolute difference in counts between the approach and the first expert divided by the number of counts measured by the first expert and then multiplied by 100) and the Dice similarity coefficient (a pixel-based measure of similarity) between each automated result and the reference were computed for each image and averaged across all images. In addition, the Pearson correlation coefficient (R) between the counts provided by each automated approach and the manual reference counts were computed. The axon counts and complete tracings were also obtained from a second expert and compared to those from the first expert (again, measuring the average absolute count difference as a percentage, the Pearson correlation coefficient of the counts, and the Dice similarity metric of the complete tracings).

In addition, to further evaluate the semi-supervised approach against a similar structured FCN, we compared these two approaches with different sizes of training data size. Both approaches were trained on different sizes of randomly chosen labeled images: 100% (26 images), 50% (13 images), 25% (6 images), 10% (2 images). Meanwhile, for each training data size, the semi-supervised learning approach was also trained on all the unlabeled/unmarked data (50 images). Metrics of absolute count difference, correlations, and Dice coefficients were then compared between the two models trained on each data size.

## Results

Overall quantitative results are summarized in Table 1 (with results separated by damage grade available in S. Table 1), with the semi-supervised approach having the best overall performance. Comparing the counts resulting from the semi-supervised learning approach with the reference counts resulted in a mean absolute percent difference in counts of 4.4%, with a Pearson’s correlation coefficient of R=0.97. The Dice coefficient (a measure of the relative overlap of the pixel-based segmentations) was 0.81 for the semi-supervised approach. In contrast, the mean absolute percent difference in counts for the AxonJ approach was significantly (*p* < 0.05 using paired *t*-test) higher (9.0%), the Pearson’s correlation coefficient was significantly (*p* < 0.05 using Williams’ test^63^) lower (R=0.86), and the Dice coefficient was significantly (*p* < 0.05 using paired *t*-test) lower (0.70). The FCN approach and second expert had similar results that were just slightly worse than the semi-supervised approach. Visually, the semi-supervised approach also performed best, yielding qualitatively more confident and clean predictions than the FCN approach (Figure 7). Compared with the AxonJ results, fewer false positive regions were segmented.

**Figure 7.**
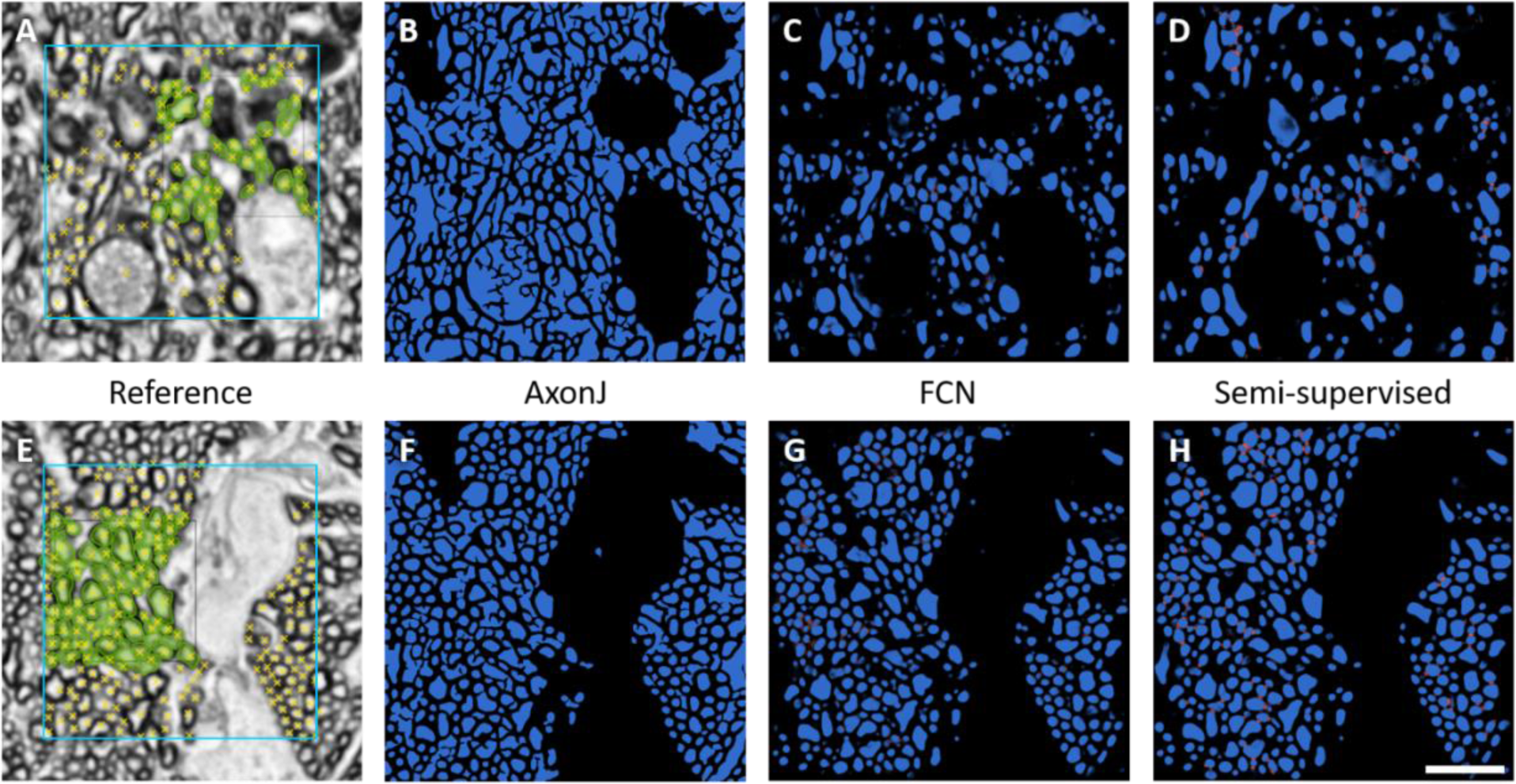
Comparing axon segmentations performed by the fully convolutional network and semi-supervised approaches relative to AxonJ and a manual reference. Two representative microscopic fields of paraphenylene diamine-stained cross sections, collected from a nerve with moderate (top row) and mild (bottom row) damage, with manual annotation (first column) and automated segmentations performed by the indicated approaches (three columns to the right). (A, E) Raw sub-fields (1024x1024 px) with manual annotation that include axon center marks (yellow x’s; inset blue border) and smaller sub-fields for axon tracings (green infilling; inset black box, 400x400 px) performed by expert #1 to provide a reference. Inset blue box denotes the border for inclusion of axons for manual center marks (larger blue border) and tracings (smaller black border). Segmentation of axons (in blue) rendered from the raw image performed by (B, F) AxonJ and the (C, G) fully convolutional network (FCN) and (D, H) semi-supervised deep-learning approaches. Red markings highlight instances in which the algorithm of each approach detected borders between adjacent axons to prevent the segmentation of multiple axons as a single axon (C-H). Scale bar = 5 µm.

**Table 1.** Comparing the performance of each approach on the final testing set.

| Metric |  | Approach |  |  |
| --- | --- | --- | --- | --- |
| (Relative to Expert #1): | Expert 2 | AxonJ | FCN | Semi-supervised |
| Abs. % diff. | 6.6 ± 5.5% | 9.0 ± 8.1% | 6.2 ± 3.6% | 4.4 ± 3.3% |
| R | 0.93 | 0.86 | 0.93 | 0.97 |
| Dice coefficient | 0.79 ± 0.04 | 0.70 ± 0.10 | 0.80 ± 0.05 | 0.81 ± 0.04 |

Footnotes. The following were abbreviated as follows: absolute percent difference (Abs. % diff.), Pearson's correlation coefficient (R), and fully convolutional network (FCN). The absolute percent difference and Dice coefficient values are reported as the mean plus/minus the standard deviation across the corresponding test images.

When visually comparing the performance of the semi-supervised approach vs FCN with different sizes of training data (Figure 8), at each training size, the semi-supervised approach showed qualitatively cleaner segmentations and less noise, with borders between axons more clearly seen compared to the FCN approach. Quantitatively, with the exception of the absolute percent difference in counts when using only 10% (i.e., 2 images) of the labeled images for training, the semi-supervised approach had a better performance (Table 2). The discrepancy between Dice metric and the absolute percent error of the counts in the 10% case may have been in part due to the counts being more affected by the post-processing stage in its removal of smaller objects.

**Figure 8.**
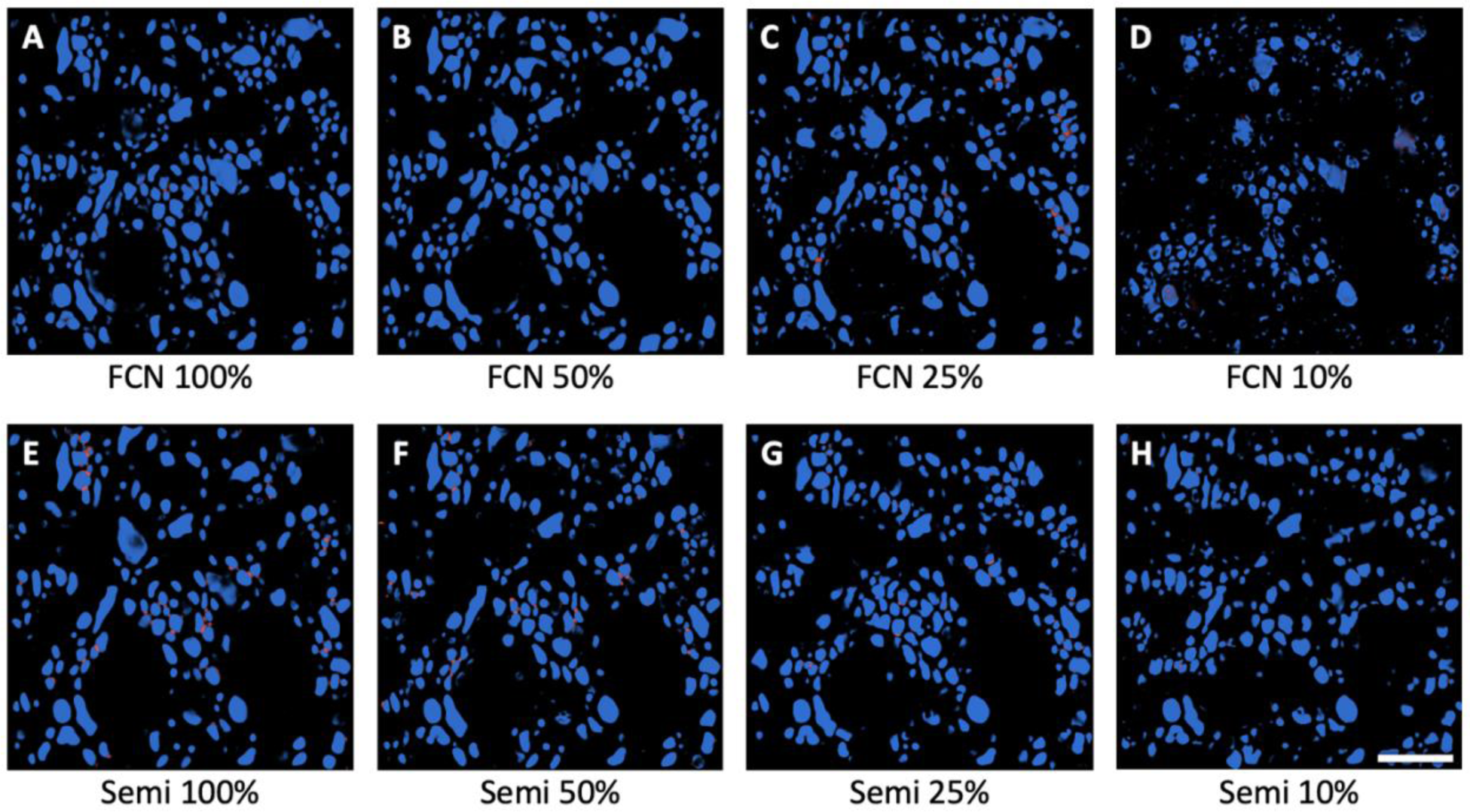
Comparing the resultant axon segmentations generated by the fully convolutional network and semi-supervised learning approaches with incremental decreases in the amount of training data. Axon segmentations (in blue) generated under direct training by (A-D) the fully convolutional network (FCN) and (E-H) semi-supervised learning approach from the same microscopic field. Non-axons are represented in black and red markings highlight instances in which the algorithm of each approach detected borders between adjacent axons to prevent neighboring axons from being segmented as a single axon (A-H). Scale bar = 5 µm.

**Table 2.** Comparison of FCN and semi-supervised approach when training on different training-set sizes.

|  | FCN<br>100% | FCN<br>50% | FCN<br>25% | FCN<br>10% | Semi<br>100% | Semi<br>50% | Semi<br>25% | Semi<br>10% |
| --- | --- | --- | --- | --- | --- | --- | --- | --- |
| Abs. % diff. | 6.2% | 5.6% | 8.1% | 9.0% | 4.4% | **4.3%** | 6.2% | 9.4% |
| R | 0.93 | 0.93 | 0.87 | 0.82 | **0.97** | 0.96 | 0.90 | 0.83 |
| Dice coeff. | 0.80 | 0.79 | 0.78 | 0.57 | **0.81** | 0.80 | 0.78 | 0.71 |

Footnotes. The following were abbreviated as follows: absolute percent difference (Abs. % diff.), Pearson's correlation coefficient (R), fully convolutional network (FCN), semi-supervised (Semi) approach, and Dice coefficient (Dice coeff.). The absolute percent difference and Dice coefficient values are reported as the mean across the corresponding test images.

## Discussion & Conclusion

Overall, in this work, we proposed a deep fully convolutional network named AxonDeep for segmenting axons in PPD-stained optic nerve cross sections from mice and presented results of training this network using a supervised approach (using only labeled data) and a semi-supervised approach (extending the network in a generative-adversarial-network framework for purposes of training with both labeled and unlabeled data). Our results show a significant improvement over AxonJ using the semi-supervised learning framework both visually and based on quantitative metrics comparing axon counts and axon segmentations with an expert reference. In the final test set, we did not include cases of severely damaged nerves as it was already known that AxonJ would perform poorly in such cases. While severe nerves were not included as part of supervised training and evaluation, here we include a few qualitative examples to indicate the increased ability of the semi-supervised approach (AxonDeep) in segmenting axons of damaged nerves versus AxonJ. For example, Figure 9 (B)(E) shows example results of AxonJ in cases of severely damaged nerves. In these cases, there are around 500 axons counted with AxonJ, compared to less than 100 axons annotated by an expert. The output from the semi-supervised learning approach can be seen in Figure 9(C)(F). Thus, the performance of this iteration of AxonDeep to recognize axons from some severely damaged nerves appears promising, but it is important to reiterate the caveat that AxonDeep has only been validated for use on nerve images with normal to moderate degrees of damage.

**Figure 9.**
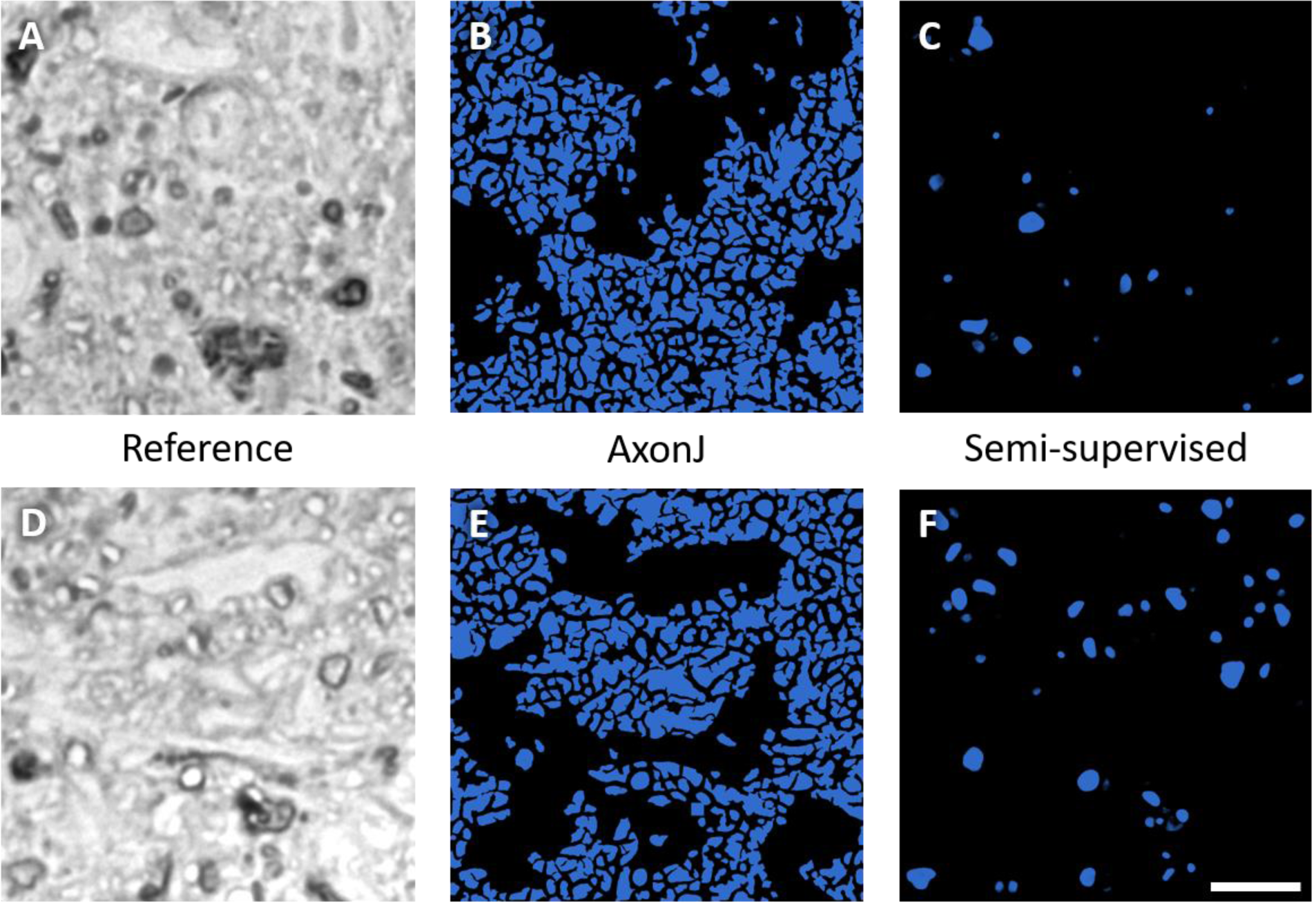
Comparison of axon segmentations performed by AxonJ and semi-supervised learning approach in optic nerves with severe damage. (A, D) Microscopic fields taken from two different optic nerves (top versus bottom row) exhibiting severe damage and corresponding axon segmentations (in blue; non-axons in black) generated by (B, E) AxonJ and the (C, F) semi-supervised learning approach. Scale bar = 5 µm.

Different from existing deep-learning approaches for providing axon counts, such as AxoNet,^28^ we directly segment axons. Use of the semi-supervised training approach to be able to still utilize untraced images was one of our strategies for more effectively dealing with the challenge and time-consuming nature of obtaining complete reference segmentations, as is needed for training. Another strategy that we used (for training purposes only) to help address the difficulty in obtaining complete reference segmentations, as previously described in more detail in the methods, was to develop an approach for editing existing segmentations rather than tracing completely from scratch. Before editing, to help avoid any bias in the number of axons marked by the existing segmentation, we only provided the starting automated result for axons whose center points were independently marked by an expert. Thus, while the boundaries of axons themselves for purposes of training may have had a bias towards the starting automated approach, the number of axons in the complete reference segmentations matched the counts provided manually and thus were not biased by the automated approach. (In a prior iteration on a different dataset, we considered editing the automated result directly without this step, but decided it had the potential for biasing the results towards the automated result too much.) As previously mentioned, while we felt this was an acceptable compromise for training, we did not want to have reference segmentations biased towards an automated approach for purposes of the final evaluation and thus obtained complete tracings on smaller sub-fields. In future work, the current version of AxonDeep could potentially be used for editing additional segmentations for training purposes.

Being able to consistently define the actual boundaries of axons is an area that could potentially be further explored in future work. For example, we noted that in both FCN and semi-supervised approaches, the Dice coefficient between the reference and semi-automatically generated axon segmentations was around 0.8-0.9 in the validation set, which was higher than the Dice coefficient of 0.79 between two experts in the test set. Despite a high degree of agreement between experts for assigning axon centers, there are instances of disagreement (Figure 10). Note the inconsistency between experts are not fully shown in axon count evaluations, since only the consistency of counts is evaluated. In the future, approaches like probabilistic U-Net could be used to help model such uncertainties.

**Figure 10.**
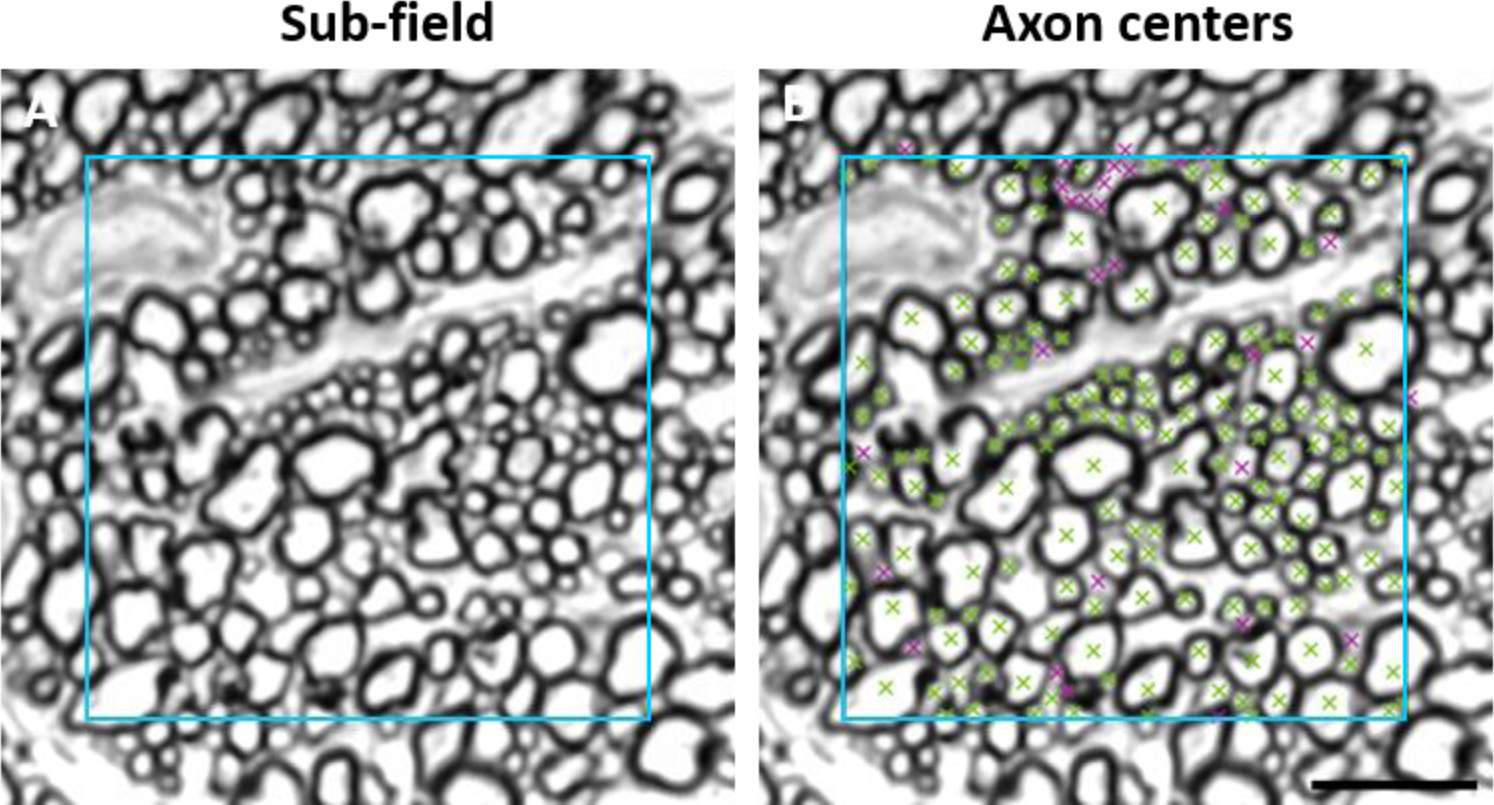
Inter-expert congruency in defining axon centers. An example of an image sub-field (A) before and (B) after manual annotation of axon center marks (colored x’s) to show inter-expert congruency. Center marks in green (green x’s) denote axons marked by both experts, whereas those in purple (purple x’s) denote axons marked by only one of the two experts (but not both experts). Inset blu box denotes the border for inclusion of axons for counting, tracing, and elimination of edge effects. Scale bar = 5 µm.

The current approach has also only been evaluated on subfields of an entire nerve. While we are currently working on an approach for quantifying axons from a montage of the entire optic nerve, we currently recommend that labs using this approach sample as many non-overlapping fields as possible, and using a separate measurement of the total optic nerve cross sectional area, mathematically convert the sum of the sampled areas into a calculated total axon number. In contexts where non-homogeneous axon densities may be suspected, we recommend quantifying successive slices of the same optic nerve (which should have near identical axon numbers), or performing the analysis twice on each section, with the microscope slide rotated by 90 degrees in the second iteration, such that averaging can be used to better control for sampling variabilities.

Implementation of AxonDeep could assist in the execution of a broad range of experiments. The mouse optic nerve contains ∼50,000 axons, though that number can vary widely according to genetic background.^45^ Quantifications of axon number is a gold-standard for measuring disease severity,^17, 64–67^ but the labor-intensive nature of manual axon counting often results in studies instead using qualitative grading scales.^19, 68, 69^ As with all of the tools that the field has put forward to count RGC somas^26, 27, 40, 70–74^ or axons^26–29, 75, 76^ in various animal models, the automated counts performed by AxonDeep greatly reduce the labor of manual counts and eliminate the possibility of user-to-user, lab-to-lab, or model-to-model variability inherent to subjective grading scales. An advantage of AxonDeep is that it performs axon segmentations as well as counts. Thus, it will also be possible to study whether axon size and shape vary during disease state. Given the high interest that has existed for many years in studying differential susceptibility of RGCs with comparatively large versus small soma,^77^ and the interesting energetic differences in large versus small axons,^78^ AxonDeep could be utilized to study many quantitative aspects of axon morphology which were previously not practical.

## Acknowledgements

We acknowledge use of the University of Iowa Central Microscopy Research Facility (a core resource supported by the Vice President for Research & Economic Development, the Holden Comprehensive Cancer Center, and the Roy J. and Lucille A. Carver College of Medicine) and the University of Iowa Multidisciplinary Investigations in Visual Science service cores (a resource supported by a P30 grant from the NIH/NEI to the University of Iowa). The contents do not represent the views of the U.S. Department of Veterans Affairs or the U.S. Government.

---

**S. Figure 1.**
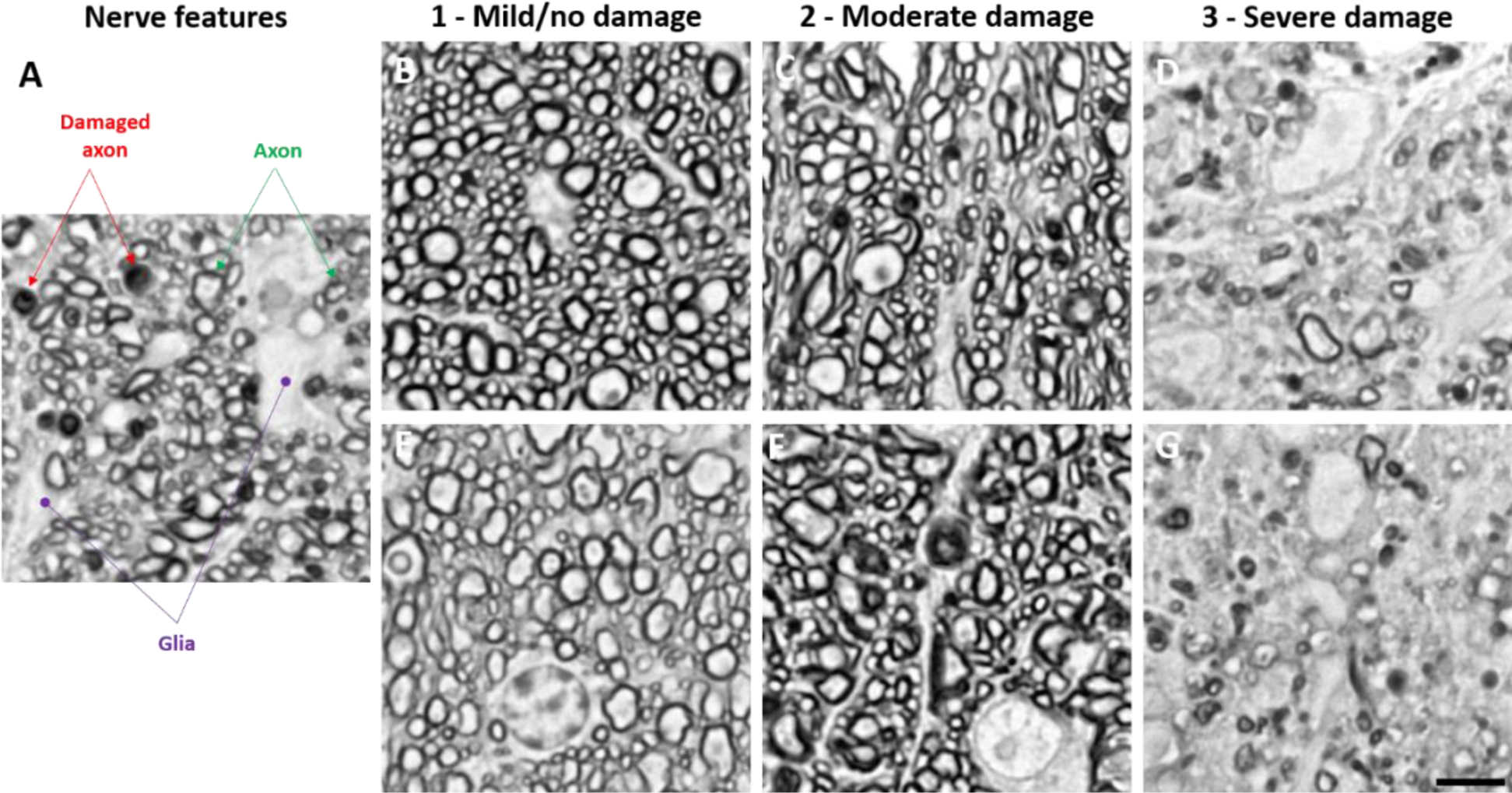
Representative images of optic nerve damage grades. Representative micrographs collected from optic nerve cross sections stained with paraphenylenediamine (PPD), a lipophilic dye which stains the myelin sheath of axons and the axoplasm of damaged axons, for each of the damage grades arranged in order of increasing damage from left to right. (A) An annotated image with examples of common features observed in optic nerves with increasing damage. (B, E) Grade 1 nerves exhibit mild to no overt damage, characterized by a dense arrangement of axons with PPD-staining limited to their myelin sheathes with an occasional instance of axoplasmic staining. (C, F) Grade 2 nerves have a moderate degree of damage, characterized by an increase of axoplasmic staining and prominence of glia. (D, G) Grade 3 nerves exhibit severe damage, characterized by a reduction of recognizable axons, increase in axoplasmic staining, and fibrosis. Scale bar = 5 µm.

**S. Table 1.** Performance of each method on nerves segregated by damage grade in the final testing set.

| Nerve damage grade | Metric<br>(relative to expert #1) | Expert # 2 | Approach |  |  |
| --- | --- | --- | --- | --- | --- |
|  |  |  | AxonJ | FCN | Semi-supervised |
| Mild/no damage | Abs. % diff. | 5.4 ± 5.9% | 7.4 ± 6.1% | 5.2 ± 3.4% | 3.5 ± 2.1% |
|  | R | 0.97 | 0.93 | 0.96 | 0.98 |
|  | Dice coeff. | 0.81 ± 0.03 | 0.74 ± 0.08 | 0.82 ± 0.04 | 0.83 ± 0.03 |
| Moderate damage | Abs. % diff. | 9.2 ± 4.0% | 12.1 ± 11.2% | 8.3 ± 3.1% | 6.2 ± 4.6% |
|  | R | 0.66 | 0.41 | 0.77 | 0.89 |
|  | Dice coeff. | 0.75 ± 0.03 | 0.61 ± 0.09 | 0.75 ± 0.04 | 0.77 ± 0.04 |
| Severe damage | * Images from nerves with severe damage were not used in test set. |  |  |  |  |

Footnotes. The following terms were abbreviated as follows: absolute percent difference (Abs. % diff.), Dice coefficient (Dice coeff.), Pearson's correlation coefficient (R), and fully convolutional network (FCN). The absolute percent difference and Dice coefficient values are reported as the mean plus/minus the standard deviation across the corresponding test images.

